# Defined cell types in superior colliculus make distinct contributions to prey capture behavior in the mouse

**DOI:** 10.1101/626622

**Authors:** Jennifer L. Hoy, Hannah I. Bishop, Cristopher M. Niell

**Author notes:** Correspondence: Jennifer Hoy, Cristopher Niell.

## Abstract

The superior colliculus (SC) mediates rapid orienting to visual stimuli across species. To determine the specific circuits within the SC that drive orienting and approach behavior toward appetitive stimuli, we explored the role of three genetically defined cell types in mediating prey capture in mice. Chemogenetic inactivation of two classically defined cell types, the wide-field (WF) and narrow-field (NF) vertical neurons, revealed that they are involved in distinct aspects of prey capture. WF neurons were required for rapid prey detection and distant approach initiation, whereas NF neurons were required for continuous and accurate orienting during pursuit. In contrast, prey capture did not require parvalbumin-expressing (PV) neurons that have previously been implicated in fear responses. The visual coding of WF and NF cells in the awake mouse and their projection targets were consistent with their roles in prey detection versus pursuit. Thus, our studies link specific neural circuit connectivity and function with stimulus detection and orienting behavior, providing insight into visuomotor and attentional mechanisms mediated by superior colliculus.

**Highlights:**

- This study provides the first demonstration of the role of specific cell populations in the superior colliculus in orienting and approach behavior.
- A genetically targeted population of wide-field vertical neurons in the superior colliculus is required for rapid prey detection and initiation of long-distance approaches.
- A genetically targeted population of narrow-field vertical neurons is required for approach initiation, accurate targeting, and approach continuity.
- Visual response properties and projection targets of these cells are consistent with their role in prey capture, linking neural circuit connectivity and function with behavior.

## Introduction

The superior colliculus (SC) plays a highly conserved role in visual processing, and mediates visual orienting behaviors across species. This includes both overt motor orienting (Comoli et al., 2012; Dean et al., 1989; DesJardin et al., 2013) and orienting of attention (Knudsen, 2018; Krauzlis et al., 2013). Given this prominent role subserving vision for action, an important goal is to determine how specific circuits and cell types within SC transform stimuli into orienting behavior (May, 2006).

The SC is a laminated structure, with the superficial SC (sSC) receiving multiple sources of visual input, while the intermediate and deeper layers receive multimodal sensory input, project to a broad range of targets and provide motor output (May, 2006). Work in rodents and other species studying the physiological properties and visual stimulus encoding of sSC cells has advanced our detailed understanding of structure-function relationships of specific neuron types in the mammalian SC (Basso and May, 2017; Cang et al., 2018; Ito and Feldheim, 2018; Oliveira and Yonehara, 2018). In particular, the classically defined wide-field (WF) and narrow-field (NF) vertical cell types (Ramón y, Cajal S Translated by Swanson N, 1995) have distinct functional and anatomical properties that indicate they may contribute to unique aspects of early visual processing to drive natural approach behaviors. WF cells have large dendritic arbors and respond to small stimuli anywhere within a large region of the visual field, making them ideal for stimulus detection. On the other hand, NF cells have narrow dendritic arbors, are direction selective and respond to stimuli within much more restricted regions of visual space, making them ideal for encoding precise changes in stimulus location. Furthermore, recent genetic studies in the mouse have demonstrated that the WF and NF cells can be independently genetically accessed via the Ntsr1-GN209-Cre and GRP-KH288-Cre lines, respectively (Gale and Murphy, 2014). However, the role of either cell type during natural visual behavior is unknown.

In previous work, we demonstrated that mice use vision to detect, orient towards, and capture live crickets (Hoy et al., 2016). The prey capture paradigm therefore provides an opportunity to determine how distinct cell types contribute to visually-guided orienting and approach behavior in a natural context. Indeed, genetic studies have revealed the circuits mediating prey capture in the zebrafish optic tectum (Semmelhack et al., 2014). However, no study to our knowledge has examined the role of specific cells types in mammalian sSC as they relate to the complex sensory-motor integration that occurs during positive orienting and approach behaviors that are critically mediated by SC (Dean et al., 1989). Instead, previous studies of the role of specific cell types in sSC have only examined innate responses to threatening visual stimuli such as an overhead looming disk (Shang et al., 2015, 2018; Wei et al., 2015). These studies showed that a population of parvalbumin-positive (PV) projection neurons was necessary and sufficient to generate behavioral responses related to detection of this stimulus. It remains unclear whether the PV neurons are uniquely engaged by looming stimuli that indicate potential threat, or whether they may be recruited to support positive approach behaviors as well.

We reasoned that determining how WF, NF, and PV cell types contribute to specific aspects of prey capture would significantly advance our understanding of cell-type and circuit-specific contributions to orienting behaviors mediated by the SC. Here we compared prey capture performance following Cre-dependent chemogenetic inactivation in either WF, NF or PV cells in sSC. Suppression of WF neurons specifically impaired prey detection behaviors while leaving orienting and pursuit behaviors intact. On the other hand, suppressing NF neurons significantly impaired orienting and pursuit behavior after approach initiation, while suppressing PV cells did not significantly impact our measures of prey capture performance. We found that these behavioral results were consistent with each cell type’s visual response properties in the awake mouse, as well as output connectivity. Our study is therefore the first to determine how specific cell types in sSC mediate stimulus detection and orienting responses towards natural stimuli, with potential for insight into related SC functions such as spatial attention.

## Results

### Targeted suppression of three populations of cells in superficial superior colliculus

To test the role of specific cell types in prey capture performance, we used Cre-expressing transgenic mouse lines to target the WF and NF neurons characterized by Gale and Murphy (2014), as well as the PV positive neurons that have been shown to mediate responses to threatening stimuli (Shang et al., 2015, 2018). We selectively suppressed the activity of these cells with a chemogenetic approach, employing inhibitory Designer Receptors Exclusively Activated by Designer Drugs (DREADDs; Zhu and Roth, 2014). This approach allowed us to suppress neural activity throughout the course of multiple prey capture bouts, up to several minutes, as opposed to the short timescale control provided by optogenetics. We specifically targeted the inhibitory DREADD, hM4Di (iDREADD), to Cre-positive cells in the superficial SC by virus injection. Adeno-associated virus (AAV) carrying Cre-dependent hM4Di tagged with mCherry was injected bilaterally into each of the three transgenic mouse lines 3-4 weeks prior to behavioral and physiological testing (**Figure 1A & B**). At the conclusion of each experiment, we performed histological analysis on brain sections to confirm expression of the iDREADDs throughout the SC, as indicated by mCherry fluorescence (**Figure 1A** and **Supplemental Figure 1**). Mice were excluded from subsequent analysis if we found that virus did not spread throughout both hemispheres of their SC. We quantified the density of iDREADD-positive neurons in both superficial and deep SC from Bregma −3.2 mm to Bregma −5mm in all experimental mice (see **Supplemental Figure 1A** for representative spread of virus through SC). We confirmed that expression covered the full medial-lateral extent in all three Cre lines, suggesting a similar distribution for all three cell types. All three lines had a high density of labeled cells in the superficial SC, with few cell bodies in the deeper SC (**Supplemental Figure 1B & C**).

**Figure 1.**
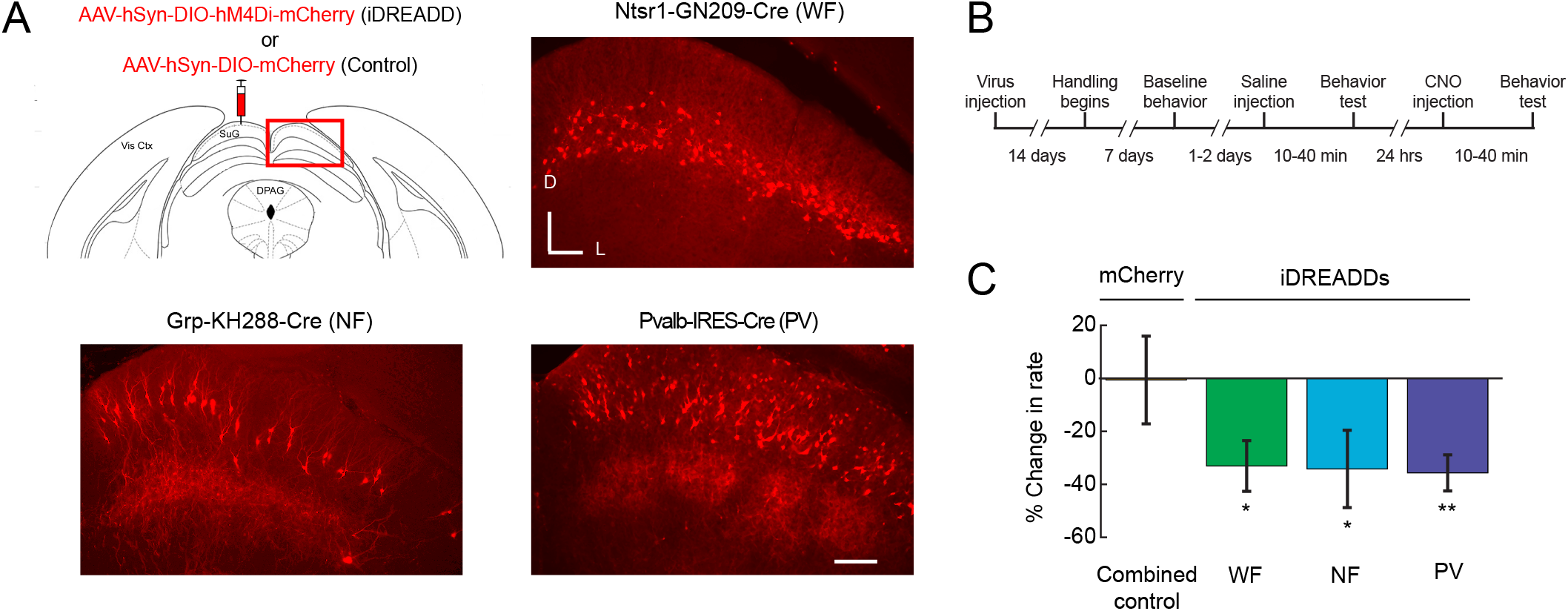
DREADD-mediated inhibition of three different cell types in mouse superior colliculus. (A) Top left: AAV virus delivering either hM4Di fused to mCherry (iDREADDs) or mCherry only was injected bilaterally into sSC, as shown in coronal section centered on the injection site (adapted from Paxinos 2004). The red box outlines the area of sSC shown in the top right and bottom panels from each Cre line injected. Top right and bottom panels: fluorescence expression pattern following iDREADD + mCherry AAV injection into three separate Cre-lines. Scale bar 100 μm, D/L = Dorsal/Lateral. (B) Timeline of experimental manipulation and behavioral assessment of each of the treatment groups. (C) Percent change in mean firing rate during stimulus presentation from before to 10 min after CNO injection, for neurons from all treatment groups. The combined control is the pooled data from all mice expressing mCherry-only in each of the three tested cell types. Control = 12 cells, 7 mice; WF = 17 cells, 5 mice; NF = 12 cells, 6 mice; PV = 18 cells, 6 mice. Significance tested using Wilcoxon signed rank test, testing for a median equal to 0, **= p<0.01, *=p<0.05. Error bars are +/− SEM.

From the histological sections, we determined the output connectivity of each cell type by measuring their axonal projections throughout the rest of the brain, to aid in interpretation of the effects of suppression on behavior. Similar to a previous report, we found that the WF cell population uniquely sent projections to lateral posterior nucleus (LP, the mouse homolog of pulvinar), whereas the NF population sent projections to intermediate and deep SC, as well as the parabigeminal nucleus (PBg; **Supplemental Figure 1D**; Gale and Murphy, 2018). The PV population projected to a number of targets, including dorsal lateral geniculate nucleus (dLGN), ventral lateral geniculate nucleus (vLGN), PBg, and patchy projections to intermediate SC, as previously reported (**Supplemental Figure 1D and Figure 1A**; Shang et al., 2015; Villalobos et al., 2018). In addition, we found a previously unreported, dense projection of PV neurons to several pretectal nuclei, including the anterior pretectal nucleus (**Supplemental Figure 1D**). Thus, the WF and NF populations have different projection targets from each other, while the PV population has a broad projection pattern that includes strong overlap with the NF targets.

To confirm that systemic injection of CNO was effective in specifically reducing the activity of Cre-dependent iDREADD-expressing neurons, we used optogenetic tagging to identify targeted cells during electrophysiological recording (Lima et al., 2009). To tag each of the cell types, we crossed the three Cre lines to the Ai32 line, which expresses ChR2 in a Cre-dependent manner (Madisen et al., 2012). 2-3 weeks prior to recording, mice from these crosses were injected in the SC with AAV expressing either Cre-dependent iDREADDs-mCherry, or Cre-dependent mCherry only as a control. Using silicon probe recordings in the SC of awake mice, we identified Cre-positive neurons based on their response to 1 ms pulses of blue light. We then measured the mean visually evoked firing rate of neurons, before and after the injection of CNO, during presentation of drifting grating stimuli. CNO injection selectively reduced the mean firing rate of all three cell types within 10 minutes (same interval as before behavioral tests) when they were expressing iDREADDs-mCherry, but not when they were expressing mCherry only (percent change = −1 ± 16%, mCherry only; −34 ± 9%, −35 ± 15 % and −37 ± 7%, iDREADD-expressing WF, NF or PV cells respectively, **Figure 1C**). The mean peak firing rate for each cell type prior to CNO injection was, 25 ± 6 sp/s, 9 ± 3 sp/s and 15 ± 6 sp/s, iDREADD-expressing WF, NF or PV cells respectively. Neighboring Cre-negative cells did not significantly change their firing rate even as Cre-positive cells were suppressed (mean percent change of Cre-negative cells in iDREADDs WF- NF- or PV- Cre mice = −8 ± 26%, −2 ± 14% and 18 ± 13%, respectively). These results confirm that CNO is effective in selectively suppressing activity of Cre-expressing neurons in each transgenic line, without significantly impacting other cell types in sSC

### Suppression of WF and NF cells differentially affects specific aspects of prey capture

In order to test the role of each of the three cell types in orienting behavior, we measured prey capture performance following iDREADD-mediated suppression via injection of CNO. All mice were habituated to the arena, handlers, and crickets after they recovered from the virus injection and before their first injections with CNO (**Figure 1B**). We compared the performance of each Cre line expressing iDREADDs in a given cell type to a single control group comprised of combined data from all three Cre lines expressing mCherry alone. All groups, including controls, received an injection of 1 mg/kg CNO 10 min prior to behavioral assessment. Thus, any potential non-specific effects of CNO would be shared across experimental and control groups.

Following CNO injection, we found significant increases in the average time to capture prey for mice with WF or NF cells suppressed by iDREADDs relative to controls (**Figure 2E**), but no effect of suppressing PV cells relative to controls. Time to capture was defined as the time between introducing the cricket and when the mouse captured and began to consume the cricket. Although suppression of WF and NF neurons increased the time to prey capture, all mice tested eventually captured and ate each cricket they were given within 7 minutes on each trial (**Figure 2, Supplemental Movies S1-4**). We confirmed that capture times for control mice injected with CNO (6.05 ± 2.70 s) were no different from mice injected with saline the day prior (9 ± 2 s, 7 ± 2 s, 5 ± 1 s, 6 ± 2 s, control, WF, NF and PV groups, respectively), demonstrating that neither the injection procedure nor CNO itself impairs prey capture performance.

**Figure 2.**
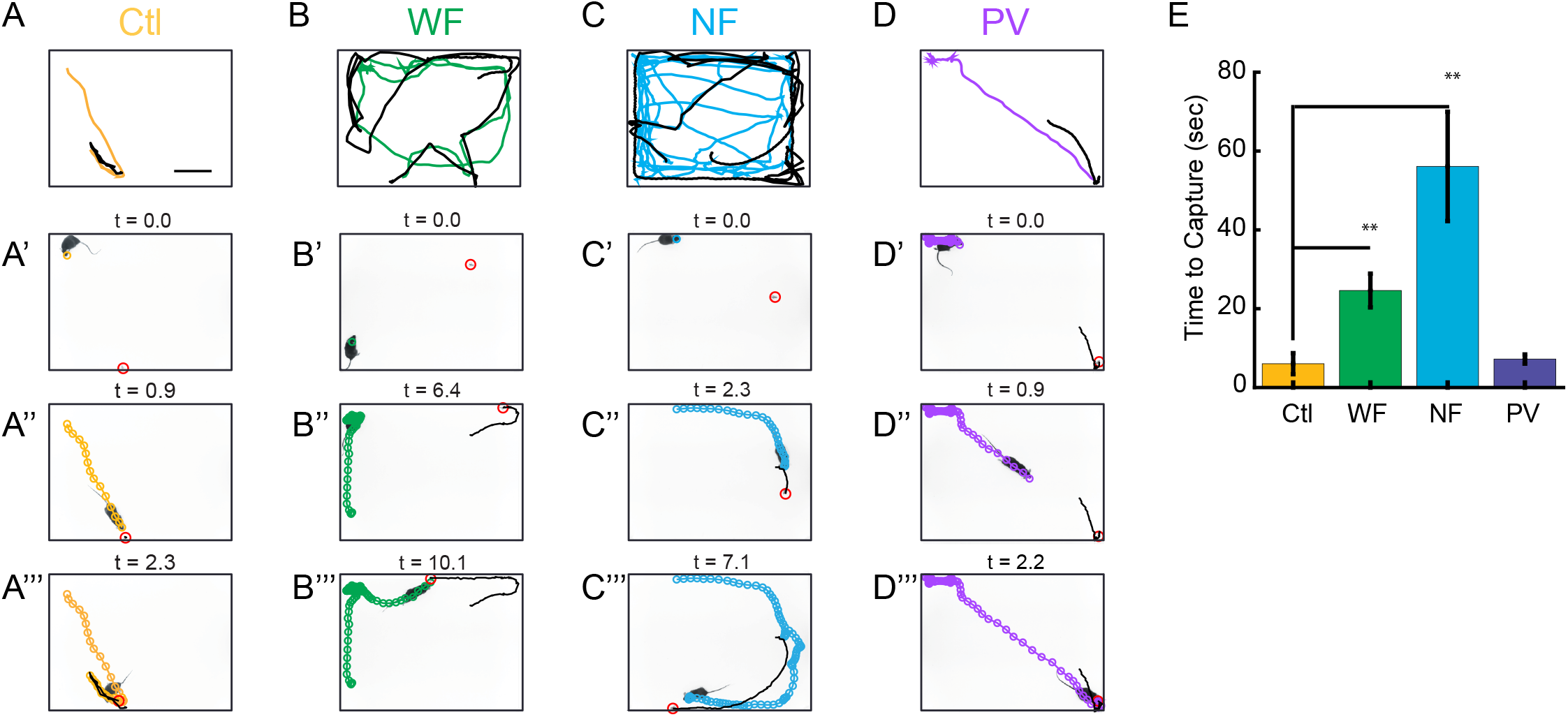
Inhibition of WF and NF cells disrupts prey capture behavior. (A-D) Top row: representative tracks from a single trial (total duration through final capture) from each experimental group, following CNO injection. Control data (mCherry only) are shown in orange, and data from mice with either WF, NF or PV cells suppressed by iDREADDs are shown in green, blue and purple respectively. Scale bar 10 cm. Bottom rows: representative detection and approach sequence (subset of timecourse shown in top row) following inhibition of cell types. (A’-A”) Control mCherry-only, see Supplemental video 1. (B’-B’”) WF cells suppressed, see Supplemental video 2. (C’-C’”) NF cells suppressed, see Supplemental video 3. (D’-D’”) PV cells suppressed (see Supplemental video 4. Note the differences in total time for each approach sequence as compared to control. Colored circle tracks denote head position marked at 16 ms intervals. Red circles highlight cricket location. (E) Median cricket capture time for each group. n=10, 10, 11,9, in control, WF, NF, PV respectively. Significance tested for by Mann-Whitney U. **=p<0.01 and *=p<0.05. Error bars are +/− standard error of the median.

To determine the distinct impairments in prey pursuit for each group of mice, we visualized individual approaches that we defined based on mouse speed and distance from target (range) relative to the cricket (**Figure 2A-D**), as in Hoy et al., 2016 (and see Methods). Briefly, approach starts were defined as moments where the bearing of the mouse relative to the cricket was within 90 deg, the range began to continuously decrease, and the mouse was moving at greater than 3 cm/s. The approach continued for as long as all behavioral criterion described above were met, or until the mouse made contact with the cricket (range < 3 cm). Under normal conditions, it takes less than 5 s from cricket introduction to when mice begin their first approach and complete a successful contact event (**Figure 2A-A’”, Movie S1**). If an initial contact does not result in capture and consumption, they quickly resume pursuit and mount another approach.

When the activity of WF cells was suppressed (**Movie S2**), mice took longer to detect the cricket, initiate their first approach (**Figure 2B”**), and started approaches closer to the target (**Figure 2B’”**, versus **Figure 2A”**), but were successful in pursuit following initiation (**Figure 2B’”**). Strikingly, when the activity of NF cells was suppressed (**Movie S3**), mice showed deficits in orienting toward the cricket, were less accurate in their approach toward the cricket, and often stopped their pursuit, becoming immobile before intercepting the cricket (**Figure 2C’-C’”**). When PV cell were inhibited (**Movie S4**), mice made the rapid, precise and distant approaches observed in control conditions (**Figure 2D-D’”** versus **Figure 2A-A’”**). Together these changes in approach and pursuit lead to significant differences in time to capture between groups (**Figure 2E**).

To more precisely quantify the deficits caused by suppression of WF or NF cell activity, we assessed distinct aspects of the behavior required for efficient prey capture (Hoy et al., 2016). In particular, we calculated parameters to assess a) detection, b) accuracy of approach, and c) continuity of pursuit.

We first quantified two measures associated with *prey detection*: time to first approach (as defined above), and range when an approach started. Suppressing the activity of either the WF or NF cells increased the average time to first approach prey relative to both controls and mice with suppressed PV neuron activity (**Figure 3A**). These results indicate that suppressing WF or NF cells impairs prey detection, as measured by the ability to initiate approaches. We note, however, that this could be due to either failure to detect the stimulus, or failure to execute the orienting that is our measure of detection. Interestingly, control mice tended to start a successful approach from either approximately 10 or 30 cm away from the target, as the approach start distribution was bimodal (see Methods for description of distribution). In control mice, 86 ± 5% of approaches were started farther than 22 cm away from the target (**Figure 3A**, bottom graph). Notably, mice with suppressed WF cells specifically showed a significant decrease in the probability of starting approaches from greater than 22 cm away from target (**Figure 3A**, bottom graph), whereas NF and PV cells showed no change in start location. Taken together, this suggests a specific role for WF cells in stimulus detection, particularly at long distances.

**Figure 3.**
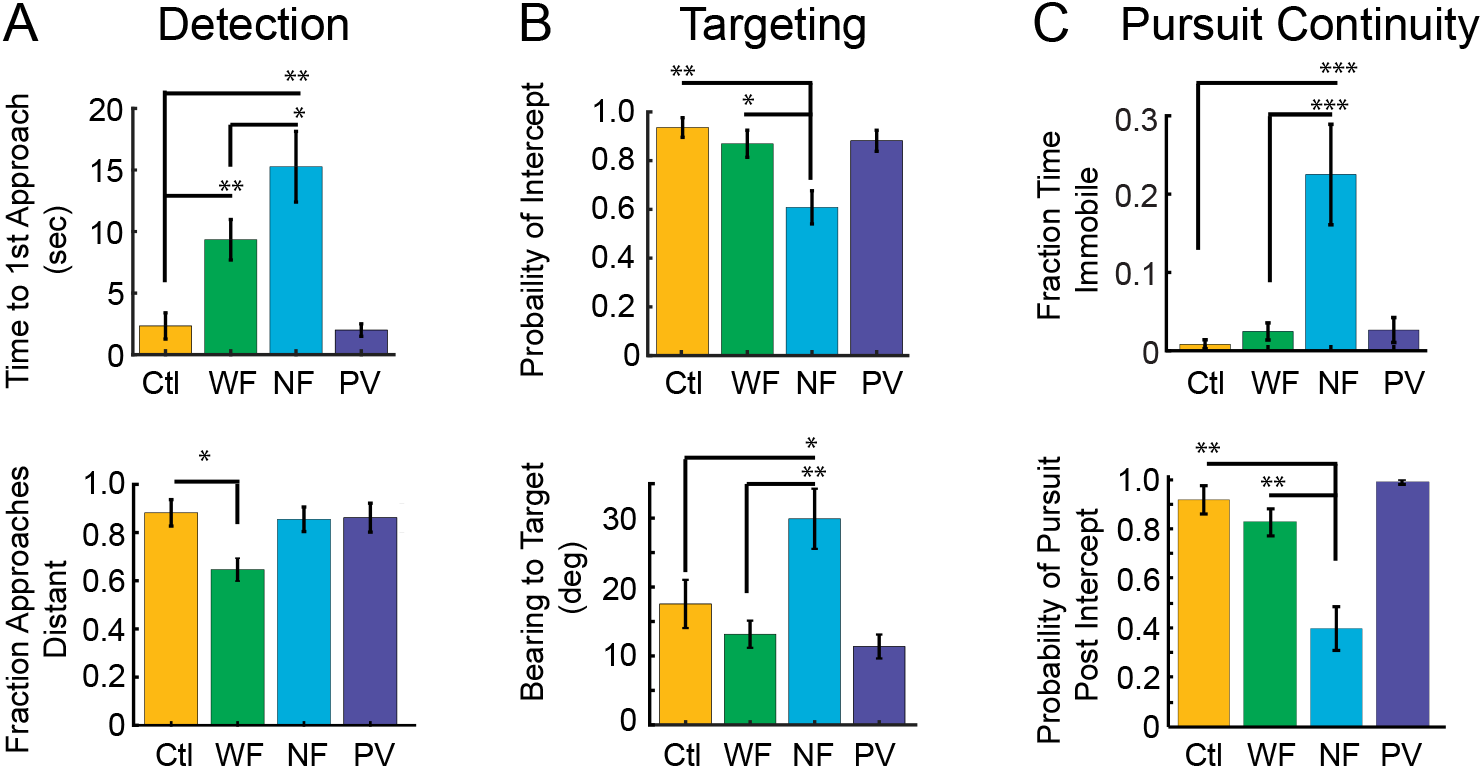
Suppression of specific cell types differentially alters target detection and pursuit behavior (A) Detection performance. Top: Median time to first approach. Bottom: fraction of approaches that begin greater than 22 cm from cricket (B) Targeting performance. Top: Mean probability of an intercept following initiation of approach. Bottom: Median bearing to target prior to an approach that ends in contact. (C) Pursuit continuity. Top: Median percent time immobile within 5 cm of the prey. Bottom: Probability mouse will successfully re-initiate pursuit immediately following interception if it fails to capture. Significance tested using Kruskal-Wallis, =0.05, followed by Mann-Whitney-U and Dunn-Sidak correction for multiple comparisons. n=10, 10, 11, 9, control, WF, NF, PV. ***=p<0.001, **=p<0.01, *=p<0.05. Error bars are standard error of the median.

Second, we quantified the *accuracy of targeting* by comparing the relative bearing to target during an approach (azimuthal angle of head relative to cricket location), and the probability of successfully intercepting the cricket given an approach. Suppressing the activity of NF cells decreased the accuracy of targeting in both of these measures, while neither WF nor PV cell suppression affected the ability to execute the approach once it had begun (**Figure 3B**). These findings suggest that NF cells in sSC play a unique and critical role in accurately targeting prey, relative to the other cell types investigated here.

Third, we quantified the ability of mice to consistently maintain pursuit of the prey (*pursuit continuity*). We observed that suppressing the activity of NF cells often resulted in mice pausing shortly before prey contact during an approach, when they were within 5 cm of the target (**Figure 3C, top** and **Figure 2C’”**, clustered track points near target position). These mice were also less likely to continue pursuit by re-initiating an approach after the cricket escaped a contact event (**Figure 3C, bottom**). Again, these differences in pursuit continuity were unique to suppressing the activity of NF cells and indicate that this population of cells is critical in driving and maintaining accurate motor behaviors during pursuit and ensuring rapid prey capture.

One possible explanation for the differences observed across the cell types could be due to differences in the number of neurons expressing iDREADDs in sSC overall. However, quantification of the density of labeled cells in each experimental group suggests that this is not the most likely explanation. A similar number of cells expressed hM4Di in the PV and WF groups, while the NF group had half as many cells expressing hM4Di (**Supplemental Figure 1B**) yet had the most severe behavioral impairment. Furthermore, our electrophysiological recordings showed a similar degree of suppression of firing rate per cell (∼35%) following CNO administration in these three groups (**Figure 1C**). Therefore, the extent of the behavioral phenotype does not correlate directly with the number of cells expressing iDREADDs nor our measures of percent reduction in stimulus evoked firing rate after CNO administration. The PV cells also overlap in the location of their cell bodies with both the WF and NF cells (**Figure 1A**), and some of their projection targets overlap with that of the NF cell population (**Figure 1A** and **Supplemental Figure 1D**). Despite sharing these anatomical properties with NF neurons, suppressing the activity of PV cells had no impact on our measures of prey capture behavior (**Figure 2**). This also suggests that the effects on behavior are cell type-specific and unlikely to be caused by regional suppression of activity in SC or other projection target areas.

To determine whether the observed phenotypes were a result of general disruption of motor behavior or other state changes such as anxiety, we analyzed running speeds and exploratory behavior in the open field in the absence of live prey after CNO injection. We did not identify any differences between experimental groups and controls in maximum and mean speeds of locomotion, time spent within the perimeter of the arena, or percent time immobile. (**Supplemental Figure 2**). None of our manipulations significantly affected the probability of capturing the cricket given an attack, suggesting that attack behaviors relying on touch or whisking were unaffected by our manipulations (probability of capture given contact was 38 ± 16%, 32 ± 4%, 20 ± 5%, and 33 ± 7%, control, WF, NF or PV mice, respectively). Taken together, these findings suggested that specific aspects of visuomotor behaviors are differentially affected by the selective suppression of these three cell types in sSC. The WF and NF neurons manipulated here are specifically required for some unique behaviors relevant to prey capture, while PV cells in sSC play little or no role under the conditions tested here.

### Visual responses of identified cell types are consistent with their role in prey capture

We next sought to determine how the visual function of each cell type relates to its role, or lack thereof, in prey capture. In a previous study, the visual stimulus response properties of both the WF and NF cells were characterized in anesthetized mouse (Gale and Murphy, 2018), whereas there has only been limited characterization of the WF and PV population response properties in the awake mouse (Bennett et al., 2019; Shang et al., 2015, 2018). In order to provide a systematic comparison across all three cell types we quantified a range of visual response properties in awake mice by performing acute extracellular single-unit recordings with silicon multisite electrodes while applying the optogenetic approach described above to identify Cre-positive neurons during recording (**Figure 4A**). We presented sparse light and dark spots of a range of diameters, both stationary or moving at a range of speeds (Piscopo et al., 2013). These stimuli enabled the quantification of spatial receptive fields via reverse correlation, ON/OFF response polarity, size and speed selectivity, and responsiveness to moving stimuli. In addition, we presented drifting sinusoidal gratings to measure orientation and direction selectivity as in Niell and Stryker (2008).

**Figure 4.**
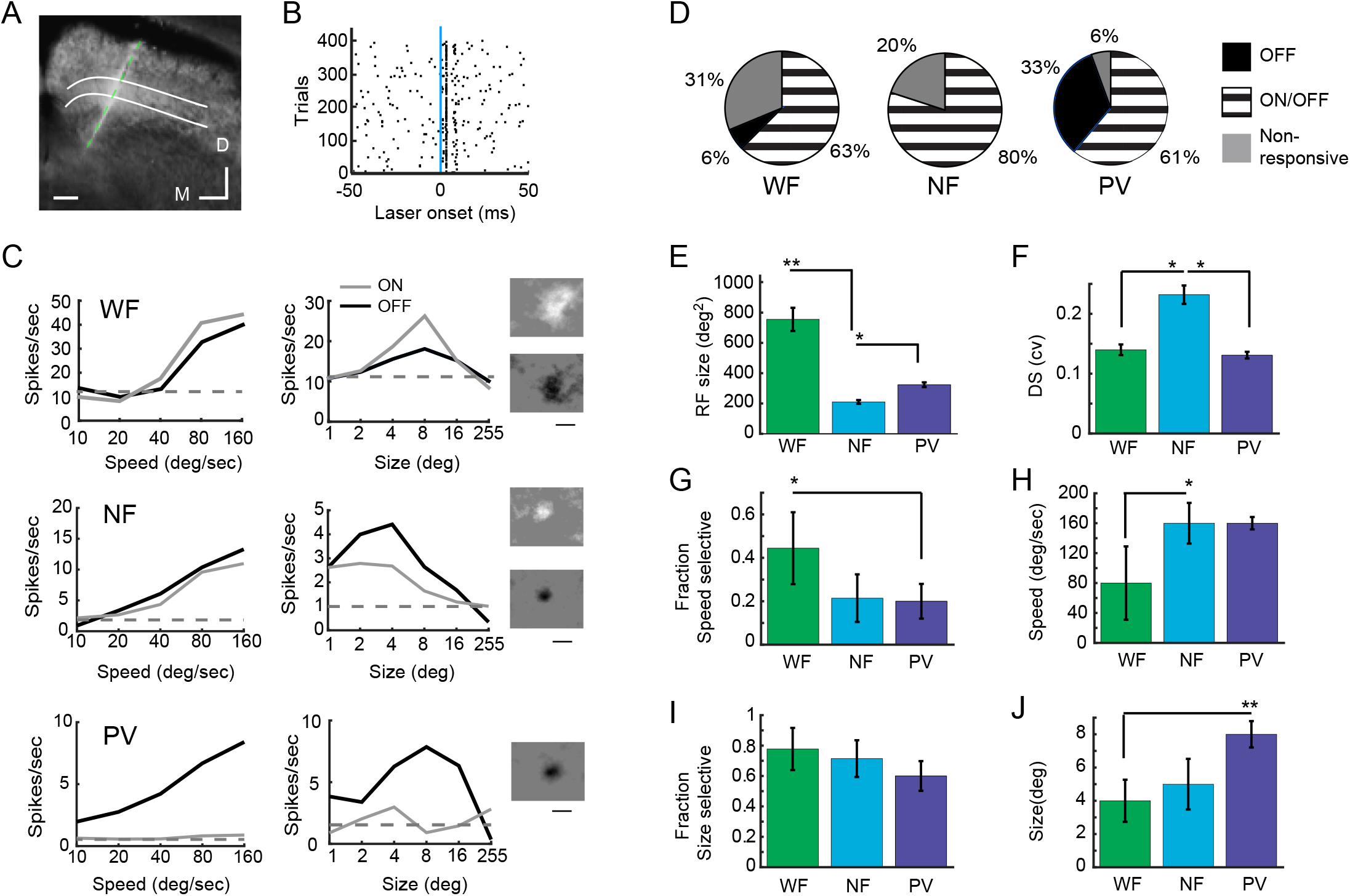
Visual response properties of WF, NF and PV cells in the awake mouse. (A) Representative image of an electrode track recovered following recording in the sSC in an Ntsr1-GN209-Cre mouse expressing Cre-dependent ChR2 in WF neurons. Scale bar equals 100 μM, D=dorsal and M=medial, parallel white bars demarcate stratum opticum. (B) Identification of Cre+/ChR2+ neurons based on response to optogenetic light stimulation. ChR2+ neurons (and hence Cre+) demonstrated short-latency responses (<1.5 ms) with little jitter (<1.5 ms) to blue light stimulation as shown by spike rasters for repeated trials. Blue line indicates laser onset time (1 ms duration). (C) Representative speed and size selective responses elicited from a WF neuron (top row), NF neuron (middle row), PV neuron (bottom row). Insets show representative spike triggered averages of the responses to moving ON (top) and OFF (bottom) spots, respectively. (D) Categorization of response types based on flashing spot stimuli, n=27, 16, and 23, WF, NF, and PV cells, respectively. (E) Estimated receptive field size based on spike triggered averages of either ON or OFF response to moving spot stimuli, for cells that were well-fit by a Gaussian function, n=14, 7, and 13, in WF, NF, and PV cells, respectively. (F) Direction selectivity by cell type. (G) Fraction of units for each cell type that were speed selective. (H) Median preferred speed for selective units, n=12, 4, and 5, WF, NF, and PV cells, respectively. (I) Fraction units for each cell type that were size selective. (J) Median preferred size for selective units, n= 21, 11, and 14, WF, NF, and PV cells, respectively. Significance tested using Kruskal-Wallis, =0.05, followed by Mann-Whitney-U and Dunn-Sidak correction for multiple comparisons. **=p<0.01, *=p<0.05. Error bars are standard error of the median. n=28, 15, and 23, WF, NF, and PV cells, respectively, except where noted. Neurons were recorded from 10, 8, and 9 Ntsr1-GN209-Cre, GRP-KH288-Cre, and Pvalb-IRES-Cre mice respectively.

Quantifying the visual response properties of the WF, NF and PV neurons to the same stimulus set confirmed that the targeted WF and NF populations have distinct response preferences from each other and from PV cells (**Figure 4C-J**). Figure 4C shows representative responses from the three cell types and demonstrates that all cells show robust responses to high speed stimuli and prefer relatively small stimuli. WF neurons have the largest receptive field sizes (**Figure 4E**), consistent with finding that they respond to small sparse stimuli anywhere within a relatively large area of visual space. They respond to both light (ON) and dark (OFF) spots and prefer moving stimuli, as they are the least responsive to flashing stimuli (**Figure 4D**). On the other hand, the NF neurons have much smaller receptive field sizes than the other two cell types (**Figure 4C & E**) and are direction selective (**Figure 4F**). While the same types of information were conveyed by subsets of the PV population, this population also had a substantial fraction of neurons with only OFF responses and a preference for larger sizes than NF and WF cells.

Overall, the receptive field properties of the WF and NF neurons mapped well to their roles in prey capture behavior. WF cells encoded small (4-5 deg) light or dark objects moving within a large (700-900 deg^2^) region of the visual field, consistent with detecting the presence of a stimulus. In contrast, the NF cells responded to small light or dark spots within a much smaller region of visual space (<200 deg^2^), and were tuned to the direction of stimulus movement, information that is important for guiding approach toward a target. The PV cells responded to larger and often dark stimuli, and while we did not find a role for the PV cells in prey capture behavior here, these response features are consistent with their previously demonstrated role in mediating fear response to looming stimuli.

## Discussion

### Specific cell types and circuits within the sSC contribute to distinct visual behaviors

Our findings connect genetically identified cell types in sSC and their visual responses in the alert mouse to specific aspects of prey capture behavior. We found that suppressing WF neurons disrupted rapid prey detection but left other aspects of prey capture unperturbed, such as accurate orienting and continuous approaches. This disruption in detection is consistent with the observation that WF neurons have large receptive fields, are selective for stimulus size, are most responsive to moving stimuli, and project exclusively to lateral posterior nucleus. The selectivity for small objects anywhere within a large receptive field could provide sensitivity for stimulus detection, with less specific information about stimulus location or direction of motion. However, it should be noted that although individual WF neurons do not encode location with high precision, a population of many neurons with significantly overlapping RFs could provide an accurate estimate of the spatial location of a small moving object (Chen et al., 2015).

Furthermore, the WF neurons project to LP, the mouse homolog of pulvinar, which could convey the detection signal to visual cortex as an “alert” for salient stimuli. Previous studies of LP have demonstrated that this higher-order thalamic nucleus encodes visual stimulus movement over a large region of the visual field, and provides information to cortex about when the movement of stimuli in the visual scene are not predicted by the animal’s own actions (Masterson et al., 2009; Roth et al., 2016). In addition, recent studies revealed that information carried by WF cells to LP is then relayed to specific areas of extrastriate visual cortex by circuitry that operates in parallel to the classical retino-geniculate pathway to cortex (Beltramo and Scanziani, 2019; Bennett et al., 2019). Therefore, our findings support the idea that WF cells in sSC provide rapid initial detection of stimulus presence and can convey this information to extrastriate cortex via higher-order thalamus.

On the other hand, we demonstrated that suppressing NF neurons impaired accurate targeting and continuous pursuit behaviors. These neurons respond best to stimuli localized within small regions of the visual field, are direction selective, and project directly to intermediate/deep SC and the parabigeminal nucleus (PBg). The small receptive fields and direction selectivity are consistent with the information needed to drive an accurate motor output to a specific location of a moving target. Moreover, the deep SC neurons are topographically organized such that they can direct eye or head movements towards a particular region in visual space by driving downstream brain regions involved in motor output (Doubell et al., 2003; Mohler and Wurtz, 1976; Schiller and Stryker, 1972). Furthermore, in many species PBg plays a role in competitive stimulus selection (Mysore and Knudsen, 2011) that may be involved in selecting one target within the visual field. Providing input to both deep SC and PBg could therefore simultaneously guide motor output during approach and facilitate selection/pursuit of a single target (Evans et al., 2018; Han et al., 2017; Li et al., 2018). Thus, our findings are consistent with NF cells driving the visually-guided motor output needed for continuous and accurate targeting of prey.

Finally, suppressing the population of cells in sSC targeted in the PV-Cre line had no significant effect on prey capture. These neurons consistently responded to large, often dark, moving stimuli, and projected to a number of targets including pretectal nuclei as well as the intermediate/deep SC and PBg. Though this population of cells is diverse morphologically, and a subset of their projection targets overlap with that of the NF population of cells we examined (Villalobos et al., 2018), our data taken together with previous studies of this cell type are consistent with the idea that these cells contribute to threat detection and associated behavioral responses (Shang et al., 2015), as opposed to prey capture. Thus, distinct circuitry in sSC is required for visual processing that drives different behavioral outputs associated with specific ethologically relevant visual stimuli.

While these results assess the contribution of WF and NF cells to prey capture behavior, it is also likely that these cells are required for other visual behaviors. We do not rule out their contribution to predator avoidance and/or novel object exploration where vision is also used to detect suddenly appearing stimuli and properly orient relative to that stimulus. In addition, we cannot completely rule out the contributions of WF and NF populations not accessed in the Cre lines selected in this study, as there is still a significant fraction of WF and NF cells in sSC that were not labeled in the Ntsr1-GN209-Cre and GRP-KH288-Cre line respectively (Gale and Murphy, 2014). Likewise, the Pvalb-IRES-Cre line labels a morphologically diverse set of neurons in sSC (Shang et al., 2018; Villalobos et al., 2018). Future studies will be required to fully address how the WF and NF cells not manipulated in our study contribute to behavior, and whether subsets of PV cells with differing morphology and projection targets differentially contribute to other visually-mediated behaviors.

Our study does not address the possible role of cortex in prey capture, as our goal was to use prey capture to probe the neural circuits for orienting and approach in the SC. However, V1 and other cortical areas provide significant input to SC that can modulate both neural activity in SC and behavioral output (Liang et al., 2015; Zhao et al., 2014; Zingg et al., 2017). It is possible that this cortical input may be required, particularly as the prey capture task becomes more complex, for example in situations that require identifying a target against a cluttered background, or fine spatial processing to discriminate appetitive from aversive small moving stimuli.

### Conserved visually-guided behaviors supported by specific SC circuitry

Though there are important species differences between mice and primates, such as the proportion of retinal input to sSC (Ellis et al., 2016; May, 2006), many visually-guided behaviors mediated by collicular circuitry are highly conserved. This is true for both rapid avoidance and orienting behaviors (King and Cowey, 1992; Van Le et al., 2013; Wang and Krauzlis, 2018). Recent studies in the mouse have identified dedicated circuitry in the SC sufficient for the initiation and completion of stereotypical avoidance behaviors, such as temporary arrest of movement in response to brief flashes of light (Liang et al., 2015; Shang et al., 2015, 2018; Wei et al., 2015), and either freezing or running towards a known “safe” space in response to overhead looming stimuli (Liang et al., 2015; Wei et al., 2015). These studies demonstrated that specific cell types in SC that responded to looming or flashing stimuli provide labeled line-like input to specific downstream targets that evoke contextually appropriate behavioral responses. Similar pathways in the SC of higher mammals are likely to exist as similar behaviors are also known to be mediated by the primate superior colliculus (DesJardin et al., 2013).

Our study suggests that cell type-specific pathways in the sSC also mediate conserved appetitive orienting behaviors. Species from across the animal kingdom naturally orient rapidly towards small, moving stimuli appearing from the peripheral visual field. The SC is required for this highly conserved behavioral response (Dean et al., 1989; Westby et al., 1990). While it is well established that the mouse sSC encodes a rich repertoire of visual stimulus features including preferences for small, moving stimuli with linear trajectories (Beltramo and Scanziani, 2019; Gale and Murphy, 2014; Inayat et al., 2015; Ito et al., 2017; Wang et al., 2010), only recently was it demonstrated that laboratory mice may use this type of visual information in the context of a specific natural behavior, insect predation (Hoy et al., 2016). We now significantly extend this understanding by defining the specific SC circuitry that is required to detect and orient towards prey (relatively small objects) from a distance. Taken together, these findings suggest that the specific neural circuits that mediate threat and reward detection discovered in the rodent may translate to the primate.

In summary, our findings significantly expand our understanding of how visual processing in the SC relates to robust, conserved ethological behavior. This opens up new directions of research studying the mouse superior colliculus to understand further aspects of approach behavior such as rapid object identification, stimulus valence assignment, target selection, behavioral choice and sustained pursuit of targets. In future work, it will be also be fruitful to further explore how the cell types accessed here contribute to other natural visually-guided behaviors in the mouse such as active predator avoidance, navigation or novel object exploration (Cooper et al., 1998; Park et al., 2018). Importantly, it is also well established that SC, particularly in higher mammals is required for more complex visuomotor and even cognitive functions, such as spatial selective attention (for recent reviews see (Basso and May, 2017; Krauzlis et al., 2013). This study begins to clarify our understanding of SC visual processing in this context by identifying specific cell types that are differentially required for aspects of visually-guided locomotor orienting. Similar processes are likely to be engaged in attentional orienting as well (Knudsen, 2018), and recent work has demonstrated that mice perform visual selective attention tasks reliably (Wang and Krauzlis, 2018). Thus, it will be interesting to determine to what extent the cell types and circuits described here play specialized roles in mediating rapid innate behaviors, or more general roles in attentional gating and salient stimulus selection across behavioral contexts.

## Supporting information

Supplemental Movie 1 - Ctrl

## Author Contributions

Conceptualization, J.L.H. and C.M.N.; Methodology and Formal Analysis J.L.H. and C.M.N.; Software, C.M.N and J.L.H; Investigation, J.L.H. and H.I.B; Visualization, J.L.H. and C.M.N.; Writing - Original Draft, J.L.H.; Writing - Review & Editing, J.L.H., C.M.N., and H.I.B.; Resources and Funding Acquisition, C.M.N.

## Acknowledgments

We are indebted to Drs. Martha Bickford, Samuel G. Solomon, Clifford Keller and Judith Eisen, and members of the Niell and Hoy labs, for helpful discussions and comments on drafts of the manuscript. We also thank Mandi Severson for extensive work obtaining and imaging histological data, and Dr. Tristan Ursell for Matlab code that facilitated color-based tracking of mouse and cricket. This work was supported by NIH grants F32EY24179 (JLH) and R01EY023337 (CMN).

## Methods

All studies were conducted with approved protocols from the University of Oregon Institutional Animal Care and Use Committees, in compliance with National Institutes of Health guidelines for the care and use of experimental animals.

### Mice

Adult male and female transgenic mice were used in this study (2-5 months of age). We used the following Cre transgenic lines to isolate and manipulate specific cell types in the superior colliculus: Grp-KH288-Cre and Ntsr1-GN209-Cre (Gerfen et al., 2013) and Pvalb-IRES-Cre (Hippenmeyer et al., 2005). In a subset of experiments each of these Cre lines were crossed to Ai32 mice in order to express channelrhodopsin-2 (ChR2) in Cre-labeled cells (Madisen et al., 2012) The mice were housed under a 12 h light/dark cycle and non-food deprived mice were provided with food and water *ad libitum*. Food trays were removed 12-24 hours before prey capture testing.

### Virus delivery

To target the expression of inhibitory DREADDs or fluorescent tags to specific populations of cells in the superior colliculus (SC), we injected AAV8-hSyn-DIO-hM4D(Gi)-mCherry (Krashes et al., 2011) or AAV8-hSyn-DIO-mCherry (gift from Bryan Roth, addgene plasmid #50459) virus bilaterally into the superficial SC of Cre-transgenic mice 2-4 weeks prior to behavioral or physiological recording experiments. 0.25 μl of virus was injected at a rate of 0.1 μl/min into each of two burr holes in each hemisphere of the skull along the following coordinates (from lambda/lateral from midline, and depth; in mm): A0.2/L0.9 and A0.0/L1.1, and D: 1.2 (Paxinos and Franklin, 2012). All mice received preoperative analgesia: 5mg/kg Carprofen (subcutaneous), lactated ringers for hydration (subcutaneous) and anesthetic: 1.5-2% Isoflurane throughout the procedure. Injections were performed with a Nanoject II (Parker, Inc.) fitted with pulled glass pipettes. Surgical sites were fully healed and sutures removed by 1-2 weeks post injection.

### Histology

Fluorescent signals were obtained from 80 μM coronal sections of 4% PFA fixed brains. Brains were imaged using a Zeiss Axio Imager 2 microscope. Images taken with the 2.5X objective were used to quantify fluorescent signals of whole structures and brain regions. To capture the mCherry fluorescent signal, we used the 1114-101 FL Filter Set 43 from Zeiss. For images, exposure times were generally set between 1000-1500 ms, with 1x analog gain.

Fluorescence images were analyzed using ImageJ analysis software. Each image was converted to 16-bit grayscale and ROIs were drawn around each of the analyzed structures: LP, dLGN, vLGN, APT, PBg and intermediate layers of the SC. Structures were identified based on (Paxinos and Franklin, 2012). We then applied the ImageJ background subtraction function and thresholded the image to select the total area containing observable fluorescent signal. We took the area fraction measure, which is the percentage of pixels in the selected ROI above threshold with significant fluorescent signal. An additional control ROI was drawn around a portion of each section outside of potential target areas in order to assess noise in this measure due to autofluorescence and other background. We quantified three representative sections approximately 160-240 μm apart through each area in each subject.

### Behavioral testing

Approximately one week prior to assessment of prey capture performance, mice were habituated to handlers, prey capture arena, and prey (house crickets, Fluker’s Farm) as described previously (Hoy et al., 2016). Experimental assessment began once mice consistently performed prey capture behavior within the arena following food deprivation, with an average capture time of less than 30 s. In order to habituate to i.p. injection, mice from all treatment groups were given an i.p. injection of saline 5 min prior to a 5 min habituation period in the arena followed by 4 prey capture trials. On the second day of saline injection, behavior was recorded and capture times for the first three prey capture trials post-injection were averaged to confirm that capture times were similar to baseline performance for each animal. The day after saline injection habituation, the same mice were all given i.p. injections of clozapine-N-oxide (CNO) at 1 mg/kg prior to arena habituation and 4 subsequent prey capture trials. All behavioral data reported here are from the first two prey capture trials, recorded within 10-40 min after CNO injection. To control for possible non-specific effects of CNO, we compared all groups to CNO injected mice from all three Cre lines expressing only Cre-dependent mCherry. Thus, our controls were virus injected and were given CNO, but expressed only the fluorophore in Cre-positive neurons in sSC.

We quantified prey capture behavioral performance as described previously (Hoy et al., 2016). Briefly, mice performed prey capture in a rectangular, white acrylic arena 45 cm long x 38 cm wide x 30 cm high with vinyl flooring. The arena was evenly illuminated from above, with luminance measured to be 60 cd/mm^2^. Video recordings were initiated prior to the introduction of the cricket. The experimental held crickets in both hands and were placed in different parts of the arena. While distracting the mouse with one hand, the insect was quietly released by the opposite hand that was not approached by the mouse. This minimized the possibility of mice employing initial localization strategies based on sound or visual cues unrelated to the prey, such as the hand of the handler. Capture time was taken as the time from the removal of the experimenter’s hands until the mouse had captured and begun to eat the cricket. The floor of the cage was cleaned between each trial and mice participated in four sequential capture trials on testing days.

Behavior was recorded at 60 frames per second with a high-resolution area scan camera (2048 x 1088 pixel; model ac2000-165umNIR, Basler). We digitized the 2-dimensional position of the cricket, the center of mass of the mouse’s body and each of the mouse’s ears (colored green with non-toxic marker or body paint) using a custom written MATLAB based algorithm courtesy of Dr. Tristan Ursell. From this tracked data, we then extracted the mouse’s head direction relative to the cricket (*θ*), target speed, mouse speed and range (distance in cm between prey and mouse). The center of the mouse’s head position was defined as the midpoint between the two ears, and head direction/bearing was defined as the vector perpendicular to the line between the two ears. From these measures we determined head position and locomotor speeds as a function of prey position over the course of each prey capture trial. All range and *θ* data were smoothed via a 50 ms sliding average window to filter out measurement noise that did not reflect the overall trajectory of the animals.

Similar to our previous studies, we defined the start of approaches as times during the capture trial when the mouse was moving at speeds greater than 3 cm/s, the range relative to the prey was steadily decreasing (Δ Range < −1cm over 50 ms), and the mouse maintained an azimuth of < 90°. We defined the termination of the approach as the moment when the mouse came into contact with the prey (defined as range < 3 cm), or when any of the initial approach criterion deviated for more than 250 ms. To control for nested data effects, we first averaged each individual’s performance over the same number of trials and then compared the mean or median of each group (except where indicated). For binominal (probability) measures, the mean (μ_p_) was equal to the probability of success (P) / number of trials (n). The standard error of the sampling distribution was computed as the standard deviation (σ = sqrt(PQ)) divided by the normalized sample size (sqrt(n)), where Q is the probability of failure, 1-P. Probability measures identified as successes in this study were: 1) if a mouse captured a cricket within a 5 min prey capture trial, p(Capture Success), 2) if a given approach towards prey ended in prey contact (end approach distance < 3 cm of prey), p(Interception |Approach), and 3) if an interception lead to the final capture, p(Capture| Interception).

### In vivo electrophysiology

Multisite silicon probe recordings in awake head-fixed mice were performed as described previously (Hoy and Niell, 2015). Briefly, mice were first implanted with a stainless steel headplate 2 days prior to recording. The headplate allowed head fixation atop a spherical treadmill (adapted from Dombeck et al., 2007; Niell and Stryker, 2010), which permits free locomotion during visual stimulus presentation and electrophysiological recording. To implant headplates, mice were anesthetized with isoflurane in oxygen (3% induction, 1.0-2% maintenance), warmed with a heating pad at 37°C and given subcutaneous injections of 0.25 ml lactated Ringer’s solution and Carprofen (5 mg/kg). In all animals, the scalp and fascia from Bregma to 2 mm behind Lambda were removed, and the skull was covered with a thin layer of cyanoacrylate (VetBond; WPI) before attaching the head plate with dental acrylic (Ortho-JET; Lang Dental Manufacturing). The well of the head plate was filled with silicone elastomer (Kwik-Sil; WPI) to protect the skull before recordings.

On the day before recording, the animals were anesthetized to perform a craniotomy over the anterior superior colliculus. The craniotomies were ∼2 mm in diameter and centered at 2 mm lateral of midline and 0.5 mm anterior of the posterior suture. The brain surface was covered in 1.5% agarose (Sigma-Aldrich) in sterile saline and then capped with silicone elastomer during an overnight recovery. After removing the protective agarose and silicone plug, the ground wire was set, and a fresh layer of 1.5% agarose in saline was applied to the well. We next inserted a 32 site, linear array optotrode (A1×32-5mm-25-177-OA32LP; Neuronexus Technologies), with 25μM site spacing and a 50μM, 0.22 NA optic fiber located ∼200μM superficial to the top electrode site. The tip of the electrode was coated with DiD (Invitrogen) to allow *post hoc* track recovery. To access the superficial SC, the optotrode penetrated overlaying V1 and was advanced until units responsive to flashing spot stimuli were observed. After the appropriate depth was reached, agarose was added to stabilize the electrode, and the preparation was allowed to settle for 40 min while the animal was shown stimuli and thresholds were set. Only one penetration per animal was used in the final analysis. All units stably isolated over the recording were included in subsequent analysis, with specific response properties calculated offline. Recording sessions typically lasted ∼2 hours and proceeded as follows: one minute light-mediated identification of Cre-dependent ChR2 expression to identify sites with ChR2 positive neurons, 20 min recording of unit responses to drifting grating stimuli, 20 min flashing spot movie stimulus, 20 min moving spot stimuli, 4 min light-mediated identification of ChR2 positive units session, a CNO or saline injection followed by 10 min settling period, and lastly, a 45 min drifting grating stimulus session to quantify post injection cellular activity. To determine the efficacy of CNO mediated shutdown in targeted cell types, the mean firing rate was calculated across all presentations of stimuli during the drifting grating presentation, pre- and post-CNO or saline administration.

After the recording session, mice were deeply anesthetized in 3% isoflurane and euthanized via cervical dislocation. The brains were then fixed whole in 4% PFA (Electron Microscopy Sciences) overnight at 4°C. Brains were subsequently sectioned and mounted in Fluoromount G with DAPI (Southern Biotechnology). We imaged the DiD electrode tracks on a Zeiss Axio Imager 2 to confirm that penetrations were specific to the target SC lamina.

### Optogenetic identification of Cre positive neurons

To identify the Cre-positive cells during electrophysiological recordings, we crossed each of our Cre lines to the Ai32 line, which expresses ChR2 in a Cre-dependent manner. This allowed optogenetic tagging of our targeted cell populations (Kravitz et al., 2013; Lima et al., 2009; Roux et al., 2014). We delivered light near the recorded neurons via an optic fiber attached to the optotrode, from a 473nm (blue light) diode-pumped solid-state laser (OEM Laser Systems, 200 mW). The presence of ChR2 positive neurons near the optotrode was first confirmed prior to recording with a brief 1 min total duration stimulation protocol whereby 1 ms long pulses of blue light were delivered at a frequency of 5 Hz for an on-period of 3 sec followed by an off-period of 3 sec. Inspecting the multiunit activity aligned to light pulse onset revealed that light-evoked neuronal activity was restricted to sites and depths consistent with the known expression of Cre-recombinase in our different lines. After recording responses to all visual stimuli, we ran the light-stimulation protocol as described above for 3 separate sessions where we varied the laser driver current (power of light illumination at the tip of the optic fiber). Light intensity at the location of the recorded neurons was estimated by measuring light power at the tip after the recording and using the calculator provided by the Deisseroth lab (https://web.stanford.edu/group/dlab/cgi-bin/graph/chart.php). Based on power emitted at the tip of the fiber at the settings used in our experiments, light intensity at recorded sites ranged from between 30-70 mW/mm^2^ with an average intensity of 41 ± 6 mW/mm^2^ at approximately 200μm from the tip. The overall duration of a session at each power was 4 min to allow for an ample number of trials to derive robust measures of reliable light-evoked neuronal responses (See Figure 4A). Units that reliably followed with a latency of less than 1.5 ms and a jitter of less than 1.5 ms across all trials after the onset of each laser light pulse were identified as putative ChR2 positive neurons and inferred to be from the targeted Cre-positive neuron population in each subject.

### Visual Stimuli

Visual stimuli were presented as described previously (Niell and Stryker, 2008). Briefly, stimuli were generated in MATLAB (MathWorks) using the Psychophysics Toolbox extension (Brainard, 1997; Pelli, 1997) and displayed with gamma correction on an LCD monitor (Planar, 30 × 50 cm, 60 Hz refresh rate). The sCreen was placed 25 cm from the mouse’s eye, subtending ∼60–75° visual space. Monitor mean luminance was measured to be 50 cd/m^2^. The monitor was centered on the hand-mapped RF location of multiunit activity.

To characterize selectivity of single-unit neural responses, we presented drifting sinusoidal gratings of 1.5 s duration at 100% contrast, with temporal frequency of 2 Hz, SF of 0.01, 0.02, 0.04, 0.08, 0.16, 0.32, and 0 cycles/° (cpd) in 12 evenly spaced directions. Gratings with a temporal frequency of 2 Hz were chosen to compare responses with those from a previous study (Gale and Murphy 2014), which showed that a temporal frequency between 1-3 Hz captures most selective responses of specific cell types in sSC. The spatial frequencies were randomly interleaved, and a gray blank condition (mean luminance) was included to estimate the spontaneous firing rate. In most cases, the spatial RF size was estimated by the spike-triggered average (STA) in response to sparse flashing noise, described in (Piscopo et al., 2013)). The sparse flashing noise movie consisted of ON (full luminance) and OFF (minimum luminance) circular spots on a gray background, at a density such that on average 15% of the area on the sCreen was covered on any given frame. Spots were 2, 4, 8,16, and 32° in diameter, and presented such that each size made up an equal fraction of the area on the sCreen (e.g., number of spots was inversely proportional to their area), to ensure even coverage at each point in space by every size. In addition, 20 frames each of full-sCreen ON and OFF were randomly interleaved. Each movie frame was presented for 250 ms followed by immediate transition to the next frame, for a total duration of 20 min. Size selectivity was estimated from the tuning curve generated from the peristimulus time histogram of response rates to each size of flashing stimuli. The size tuning was taken as the peak of the size tuning curve.

We mapped stimulus speed selectivity using a sparse moving noise stimulus (Piscopo et al., 2013). Briefly, this stimulus also uses ON and OFF spots, with a more limited size range (4, 8, 16° diameter), but each spot was assigned to move in one of eight evenly spaced directions and one of 5 speeds (10, 20, 40, 80, 160°/s). Spots appeared on the appropriate edge of the sCreen and moved across until they disappeared on the far edge. The movie was presented for 20 min total duration. Size and speed tuning curves generated for SC neurons from these stimuli were also robust, demonstrating peak responses greater than non-peak responses by at least the 1.5 Hz. In the case where a cell responded to both flashing and moving stimuli well, both stimulus sets yielded similar size tuning responses.

In some cases, subsets of WF and NF cells responded poorly to flashing stimuli (**Figure 4D**), but did respond well to sparse, moving stimuli as in (Gale and Murphy, 2014). Therefore, in a subset of experiments we used a sparse, moving stimulus to estimate the RF size and confirm stimulus size selectivity for all cell types.

### Data Acquisition, Analysis and Statistics

Signals were acquired using a System 3 workstation (Tucker Davis Technologies) and analyzed with custom routines written in MATLAB (Hoy and Niell, 2015; Niell and Stryker, 2010; Piscopo et al., 2013). For single-unit activity, the extracellular signal was filtered from 0.7 to 7 kHz and sampled at 25 kHz. Single-unit clustering and spike waveform analysis was performed as described previously (Niell and Stryker, 2008) with a combination of custom software in MATLAB and Klusta-Kwik (Harris et al., 2000). Typical recordings yielded ∼6-10 single units. Most units appeared predominantly on a single recording site, although units often contributed a signal below the voltage threshold on neighboring sites, which allowed improved unit discrimination.

For all datasets, we inspected for the presence of outliers or skews in order to determine whether to use parametric or nonparametric approaches for tests of statistical significance. For distributions that appeared bimodal, such as approach start distances, we fit distributions using MATLAB’s Statistic and Machine Learning Toolbox function as a mixture of two normals. This fitting yielded parameter estimates, means and standard deviations, for each mode. This allowed us to compute Ashman’s D, and D>2 was taken as evidence of bimodality and clear separation of modes. To determine the threshold for categorizing an event as belonging to one mode or the other, we determined if it was less than the first peak +1 SD (part of the first mode) or greater than the second peak −1 SD (part of the second mode). In the case of the approach starts distribution, the mean estimate of the first peak was 9.6 with a SD of 1.5 cm, while the mean estimate of the second peak was 31.8 with an SD of 9.4cm. Distant approaches were therefore defined as starting from greater than 22.4 cm from target. To compare means of unimodal, normal distributions, we used one-way (genotype) ANOVA followed by Tukey-Kramer HSD post hoc testing. To detect significant differences between medians, we used Kruskal-Wallis omnibus testing, followed by pairwise Mann-Whitney U tests. In some cases, we employed the Wilcoxon signed rank test to determine whether measurements significantly deviated from 0 in each tested group. Alpha in all cases was set to 0.05 and significance was inferred if p values were less than 0.05, indicated by asterisks in figure legends. Standard Error of the Median (SEM) was obtained via a bootstrapping procedure (data sampled 1000x). For all experiments we report number of trials or cells (n) as well as the total number of independent animals in each experiment in the figure legends. Unless otherwise noted, we control for nested effects by computing statistical analysis on average measures from each animal. In all cases, we also minimized nested effects by obtaining similar number of cells or trials from each animal. We typically obtained 2-4 cells or behavioral trials from each animal. Differences in frequency of categorical observations between groups were determined via X^2^.

## Supplemental Videos

**Movie S1 -** Representative prey capture trial in a control mouse (expressing mCherry only) following CNO injection.

**Movie S2 -** Representative prey capture trial during suppression of WF neurons following CNO injection.

**Movie S3 -** Representative prey capture trial during suppression of NF neurons following CNO injection.

**Movie S4 -** Representative prey capture trial during suppression of PV neurons following CNO injection.

---

**Supplemental Figure 1.**
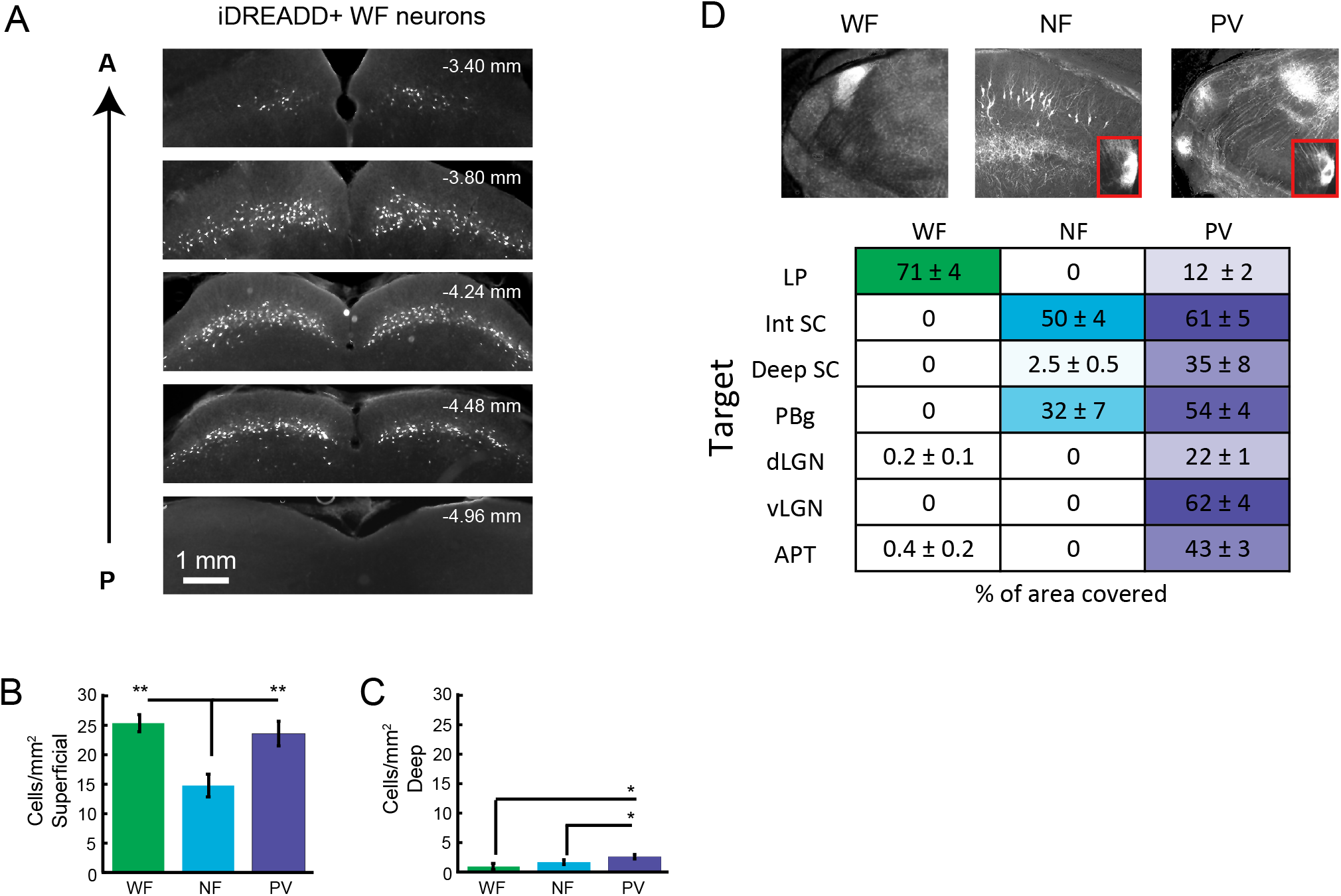
Histological characterization of expression and projection patterns in three Cre lines. (A) Representative distribution of mCherry expression throughout sSC, from an Ntsr1-GN209-Cre mouse injected with AAV-hSyn-DIO-hM4Di-mCherry (iDREADDs). A = Anterior and P = Posterior. (B, C) Mean density of each cell type expressing iDREADDs (as indicated by mCherry tag) from mice used in behavioral experiments for superficial (B) and intermediate/deep (C) layers of SC. (D) Representative images of axonal projections of the iDREADDs expressing cells in each Cre line. The coronal section with the densest projections for each line is shown, insets show PBg projections. (E) Quantification of axonal projection density to different target regions for each Cre line. We quantified fluorescence in the following structures: Lateral Posterior Nucleus (LP), dorsal Lateral Geniculate Nucleus (dLGN), ventral Lateral Geniculate Nucleus (vLGN), Anterior Pretectal Nucleus (APT), Intermediate Superior Colliculus (Int. SC), Deep Superior Colliculus (Deep SC) and Parabigeminal Nucleus (PBg).

**Supplemental Figure 2.**
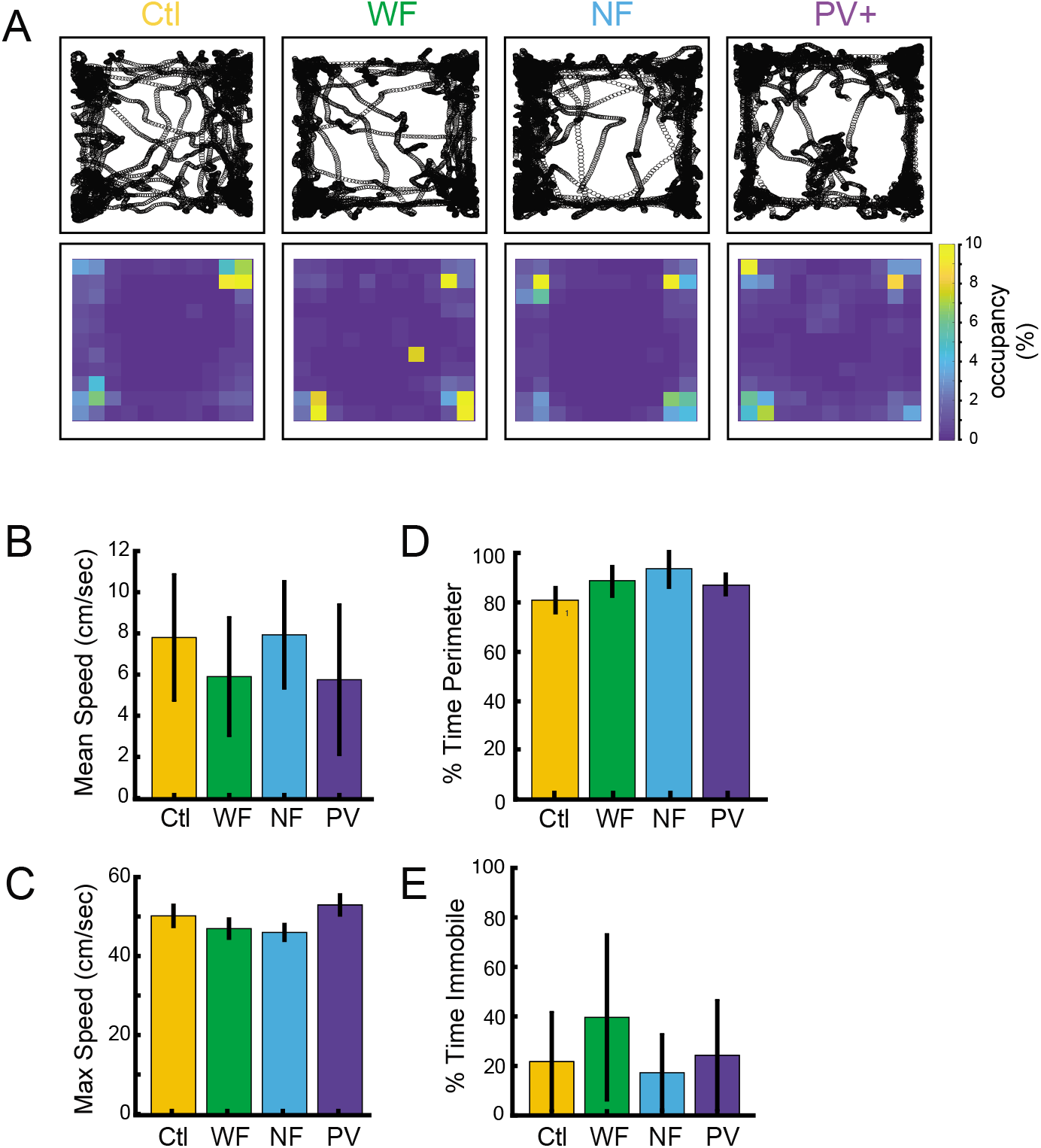
Spontaneous locomotion and exploratory behavior are unaffected by iDREADD-mediated suppression. (A) Representative tracks (top) and occupancy maps (bottom) for all four groups of experimental animals during free exploration of the arena in the absence of prey following CNO injection. (B) Mean and (C) maximum locomotion speed during free exploration following CNO injection. (D) Mean percent time spent in the perimeter of the arena and (E) percent time immobile following CNO injection. n= 20, 22, 26 and 20 trials, number of mice = 10, 10, 11, 9, in controls, WF, NV and PV respectively. Significance tested by Kruskal-Wallis, =0.05. Error bars are +/− SEM. Ctl= mice from all three lines with Cre positive cells expressing mCherry only and injected with CNO.

